# Relieving the transfusion tissue traffic jam: a network model of radial transport in conifer needles

**DOI:** 10.1101/2024.04.19.590137

**Authors:** Melissa H. Mai, Chen Gao, Peter A. R. Bork, N. Michele Holbrook, Alexander Schulz, Tomas Bohr

## Abstract

The linear geometry of conifer leaves (e.g., pine needles) imposes architectural constraints on solute transport. The needle’s structural solution to prevent axial stagnation, however, introduces an additional challenge to radial transport by restricting loading and unloading of sugar and water, respectively, to a narrow zone at the periphery of the vascular bundle. Moreover, a Casparian strip blocks apoplastic flow through the endodermis between the vasculature and photosynthetic tissue, forcing countercurrents of water and sugar to travel simultaneously through the cell lumen at this interface. In between these two potential bottlenecks is the transfusion tissue, a distinctive anatomical feature of conifer needles. Here we develop a network-based mathematical model to explore how the structure of the intervening transfusion tissue facilitates radial transport of sugar and water. To describe extravascular transport with cellular resolution, we construct networks from images of *Pinus pinea* needles obtained through X-ray μCT, as well as fluorescence and electron microscopy. Our results show that the physical separation of sugar and water pathways within the transfusion tissue mitigates the consequences of constricting flow at both the vascular access points and the endodermis.

**SIGNIFICANCE:** The efficiency with which plants transport water and sugar across many scales affects their survival and success. At the leaf scale, accommodating the opposing flows of water and sugar is a nontrivial challenge, especially when foliar geometry imposes architectural limitations (e.g., in linear leaves (needles) of conifers). The transfusion tissue is a characteristic component of the conifer needle which mediates the radial transport of sugar and water. Our network model shows that the transfusion tissue alleviates the sugar-water traffic jams that would otherwise develop between the vascular system and photosynthetic cells. Our mathematical model provides finer resolution than what is currently possible with experimental approaches.

---

Plants synthesize sugars via photosynthesis, using carbon dioxide absorbed from the atmosphere. Yet, for every acquired molecule of carbon dioxide, hundreds of water molecules are lost by evaporation through the stomata in a process known as transpiration. To avoid dehydration, the xylem, consisting of dead, thick-walled cells, passively transports water through the plant from the soil to the atmosphere under negative pressures generated by transpiration. Sugars move through the phloem from the photosynthesizing leaves to sites of storage or growth elsewhere in the plant. Because they retain their plasma membranes, phloem cells can generate positive osmotic pressures to push the flow of sugary sap down its pressure gradient. Together, the xylem and the phloem form the plant’s vascular system, which is responsible for long-distance transport of materials through the plant’s body.

In conifers, the loading of sugars into the phloem is thought to be an entirely passive process, driven by the concentration gradient of sugar between the photosynthetic mesophyll and the phloem (1, 2). For a long, narrow needle, as is typical of many conifers (e.g. *Pinaceae*), loading of the single central vein has previously been conceptualized as a continuous tube that is simultaneously and uniformly loaded along its entire length. However, mathematically solving this model predicts that the pressure at the needle base would exceed what diffusive loading can passively generate beyond an effective length of a few centimeters upstream of the base. With this paradigm of uniform loading, for long needles, phloem transport stagnates beyond this effective distance, and sugars that are produced in the upper portions of the needle cannot be exported (3–5). That this is actually the case seems unlikely, given the global success of conifers, some of which, such as *Pinus palustris*, have needles that can exceed 40 cm in length.

Liesche et al. (2021) found that needles circumvent this axial transport problem by adopting a tiered architecture in their axial venation. Moving down the needle from tip to base, new conduits are observed at the periphery of the vascular bundle at regular intervals, consistent with observations of elongation from a meristem at the needle base (6–8). This infrastructure longitudinally segments the needle into discrete loading zones that, individually, are short enough to prevent stagnation (9). However, solving the axial transport problem via the tiered architecture of the central vein introduces a new challenge in extravascular, radial transport. Photosynthesis occurs along most of the needle’s circumference, yet loading only occurs on the outer edge, or flank, of the vascular bundle (Figure 1A). As a result, sugar has only two points of entry into the phloem (10, 11). This introduces a bottleneck within the radial transport pathway that the needle must overcome. At the same time, water must flow in the opposite direction from the xylem to irrigate the mesophyll and replace water lost to transpiration. Tracer experiments show that water also flows through focused points at the flanks of the xylem before spreading out to irrigate the entire perimeter (unpublished).

**Fig. 1.**
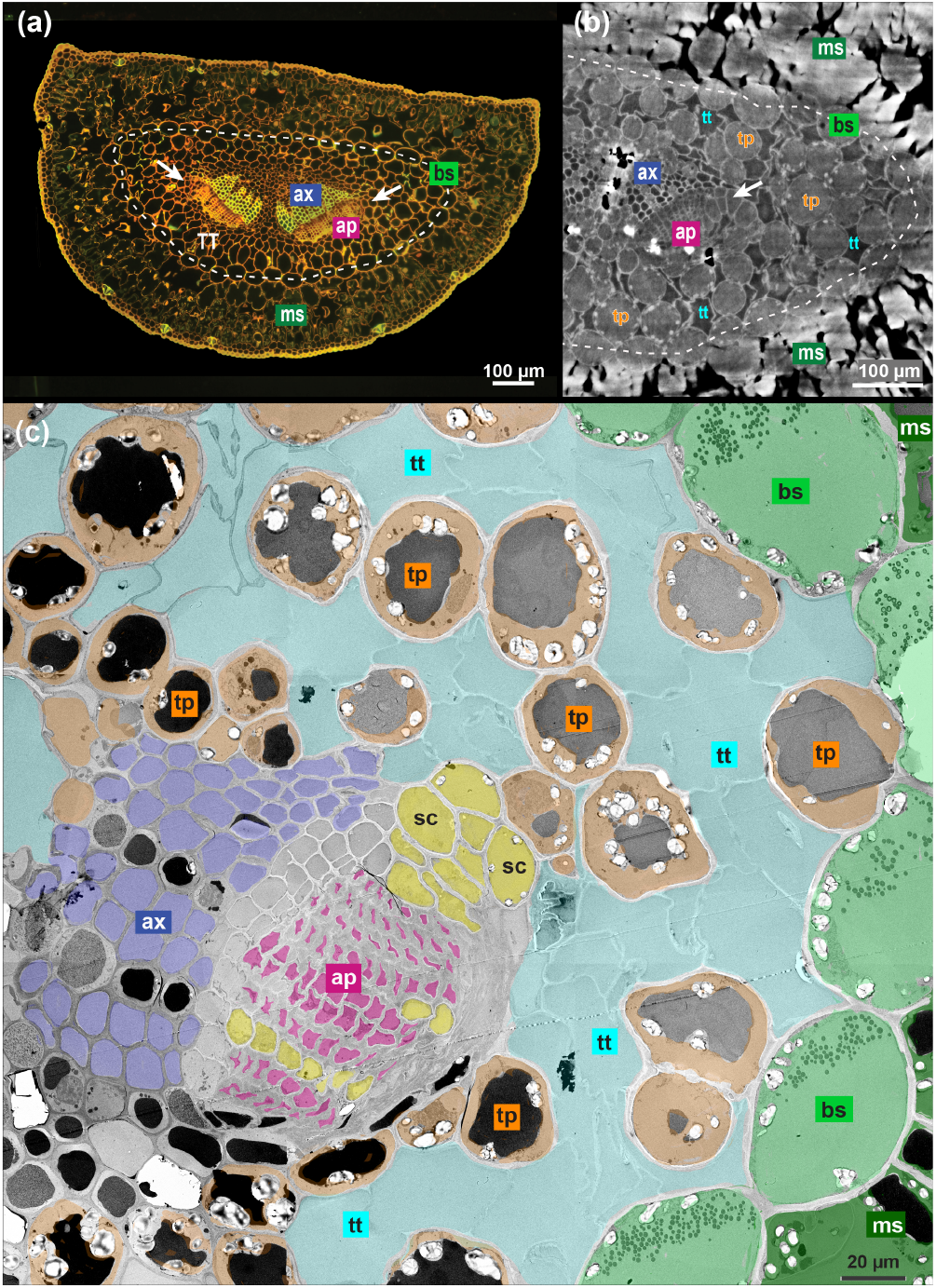
Cross-sectional architecture of a *Pinus pinea* needle as seen in fixed (a, c) and live material (b). Confocal image (a) of a coriphosphine-stained section, μX-ray computed tomograph (b), and transmission electron micrograph (c) with labels showing the tissues and cell types covered by the network model: axial xylem (ax, blue); axial phloem (ap, magenta); transfusion tissue (TT) consisting of transfusion tracheids (tt, cyan) and transfusion parenchyma (tp, orange); bundle sheath (bs, light green); and mesophyll (ms, dark green). Strasburger cells (sc, yellow) are also indicated, though they are not explicitly described in the model. Starch granules are clearly visible in the transfusion parenchyma and bundle sheath of (c). Scale bars: 100 μm (a, b), 20 μm (c). Arrows point to the flanks of the axial phloem.

Towards the exterior of the needle, transport faces another bottleneck. The endodermis serves as the gatekeeper between the vasculature and the photosynthetic mesophyll tissue. Water moves through the endodermis into the mesophyll to support photosynthesis and transpiration, while sugar moves in the opposite direction from the mesophyll into the endodermis. The endodermis consists of bundle sheath cells that are sealed with a Casparian strip, which blocks apoplastic (extracellular) flow (11, 12). By preventing flow through the cell wall, the Casparian strip forces both water and sugar to travel simultaneously through the lumen of the bundle sheath cells, despite moving in opposite directions. Here we explore how the radial infrastructure of the needle reconciles the two bottlenecks of these flows.

Characteristic of all gymnosperms is not only an un-branched leaf vascular design but also an anatomical feature known as the transfusion tissue, which lies between the axial vasculature and the endodermis (Figure 1). Fossil evidence suggests that the transfusion tissue had evolved by the late Paleozoic, with precursors appearing slightly earlier in the Permian (13, 14). Today, the transfusion tissue is ubiquitous among conifers, *Ginkgo*, and cycads, with substantial diversity in its extent and arrangement within the needle (15–20). Specifically, the transfusion tissue can adopt one of seven morphological classes, including, for example, the *Pinus*-type that encircles the entire bundle, the *Pseudotsuga*-type that adopts a U-shaped arc around the abaxial (lower, phloem-facing) side of the bundle, and the wing-like *Cupressus*-type that extends laterally from the flanks of the axial vasculature (21). Transfusion tissue is comprised of two cell types: transfusion tracheids and transfusion parenchyma. The transfusion tracheids are dead, water-conducting cells that are generally regarded as extensions of the xylem. Transfusion parenchyma are living cells that conduct sugar symplasmically to the phloem loading point, where Strasburger cells, historically labeled as *albuminous cells*, aid with loading (22).

While the structure of the transfusion tissue has been well described, its physiological function has been sparsely explored. Its physiology has been considered primarily for its role as a reversible resistor during drought (23, 24), but not for its role during transport under normal conditions. Moreover, the role of its distinctive spatial organization in the functioning of the needle has not yet been explained. The transfusion tracheid and transfusion parenchyma systems are often treated as extensions of the xylem and phloem, respectively, and were originally hypothesized to stem from the same ontogenetic origin as the primary vasculature (25). Later studies questioned the origin of the transfusion tissue in the procambium (16, 26–28). Its confinement by the endodermis, however, emphasizes that it is exclusively responsible for the two-way traffic of water and sugar between the vasculature and photosynthetic tissue (19). A striking distinction between the transfusion tissue and the primary vasculature is that the transfusion tracheid and transfusion parenchyma cells do not form spatially distinct cell files, as the xylem and phloem do. Rather, they are interdigitated within each other, with a large interfacial surface area between the two cell systems (11, 29).

Here we present a network-based mathematical model of the transfusion tissue, built to understand its role in mediating the radial two-way transport of sugar and water within the needle. As this system is difficult to probe experimentally, our computational approach resolves intercellular flows of sugar and water through the transfusion tissue, allowing us to study its hydraulic function even with uncertainty in the parameter space. We simulate networks of varying sizes and compositions to explore the impact of the transfusion tissue on transport in a functional needle. By perturbing intercellular connections within anatomically-informed networks of *Pinus*-type needles, we study how the transfusion tissue’s heterogeneous composition alleviates the constriction of flow at both the axial vasculature’s focused access points and the endodermis by physically separating the opposing currents of sugar and water.

## MATERIALS AND METHODS

### Plant material

For fluorescence and electron microscopy, *Pinus pinea* branches were collected from a 4-year-old tree in the greenhouse of the University of Copenhagen, Denmark, and, for μXCT synchrotron scanning, from a tree in the old Botanical Garden of the University of Zürich.

### Fluorescence and electron microscopy

After gently removing the epidermis, 10-mm needle segments were immediately immersed into Karnovsky’s fixative (4% [w/v] paraformaldehyde and 5% [w/v] glutaraldehyde in 0.1M sodium cacodylate buffer, pH 7.4), and fixed according to Ref. (30). To avoid preparation artefacts, 3-mm ends of each segment were discarded. After polymerization, the samples were trimmed for fluorescence and electron microscopy. Semithin sections of 2-μm thickness were transferred onto drops of distilled water placed on a microscope slide, dried on a heating plate and stained with 3% (w/v) Coriphosphine O (TCI Europe, Zwijndrecht, Belgium CAS 5409-37-0) for 5 minutes. Images were taken with a wide-field fluorescence microscope (Nikon Eclipse 80i, Amsterdam, Netherlands) at 450-490 nm excitation and 520 nm long-pass emission filters. Ultrathin sections from *P. pinea* of 70 nm thickness were cut with a diamond knife and ultramicrotome (EM UC7; Leica Microsystems, Wetzlar, Germany). The sections were transferred to film-coated single-slot grids (FCF2010-CU, Electron Microscopy Sciences; Hatfield, PA 19440, USA) and post-contrasted with uranyl-less solution for 3 min and lead citrate solution for 3 min with thorough washing after each step. For transmission electron microscopy (TEM), a ThermoFisher Talos L120C G2 was used at 120 kV acceleration voltage for an overview, stitched from 9 image tiles with 510× primary magnification using MAP3 software.

### X-ray μCT at the TOMCAT beamline, Paul Scherrer Institute, Switzerland

Needles were detached from the branch under water and immediately transferred to 2 mL Eppendorf tubes with Milli-Q water. The needles were held upright during rotation in a chamber while immersed in the Eppendorf tube (11). For imaging, the middle, tip and base segment were targeted, 1000 projections were acquired at 21 keV over 180° rotation around the needle centre using a 20× objective, resulting in an effective pixel resolution of 0.325 μm. Reconstruction of the scans followed the gridrec and paganing algorithms established at the TOMCAT beamline, providing z-stacks of 2160 needle cross sections. To avoid optical artefacts at the ends of the scanned area, 2D-segmentations were taken at cross section 500, 1000 and 1500. The dimensions of the scanned image cube were c. 0.7×0.7×0.7 mm.Nodal networks were constructed from μCT images of 12 needles, selecting cross-sections from each needle’s lower, middle, and upper thirds. Due to the needles’ bilateral symmetry, networks were generated from one half of each cross-section, retaining only sections where the endodermis was entirely visible, to create 66 total networks. Using MATLAB 2022b’s Image Processing Toolbox, cells in each image were manually identified by type. Connections between adjacent cells were then documented in an adjacency matrix, which, together with a vector of cell identities, formed the primary inputs for the mathematical model.

### Mathematical model

The model explores the potential interference of the sugar and water flows, whose net movements run in opposite directions between the axial vasculature and the photosynthetic mesophyll. While the water flow must spread out from a concentrated source at the flank of the axial xylem to irrigate the needle’s perimeter, sugar funnels in from the circumferential region of the photosynthetic mesophyll into a concentrated loading point at the flank of the axial phloem. To avoid needing to specify spatial and geometrical details of the needle, we describe the stele (i.e., the axial vasculature and transfusion tissue), the surrounding bundle sheath cells of the endodermis, and photosynthetic mesophyll as a nodal network. Though informed by anatomy, the model does not rely on the exact positions or shapes of cells but rather on a description of each cell’s neighbors. This approach is justified by the assumption that cytoplasmic streaming keeps the cytosol well mixed and that the intercellular connections of the plasmodesmata, aquaporins, or bordered pits account for most of the resistance to flow (31). Therefore, the actual transport distance becomes less important than the discrete number of cells that are traversed (32).

A diagram of the basic model is presented in Figure 2A, and Table 1 provides the relevant parameters. Each node in the network corresponds to a cell, defined by its cell type and described by its pressure and sugar concentration. To allow for the extension of the model to non-steady conditions, cellular starch content is also included, even though its inclusion does not affect the steady-state solution. The cell types include the dead tracheary elements (axial xylem (ax) and transfusion tracheid (tt)) and the living cells (axial phloem (ap), transfusion parenchyma (tp), bundle sheath (bs), and mesophyll (ms)). The edges, or connections between nodes, correspond to intercellular connections, defined by the identities of the paired cells, and parameterize the water and sugar flows between cells. Water flows through each connection (blue arrows, Figure 2A), but sugar can only flow between living cells (red arrows, Figure 2A).

**Table 1.** Model variables and parameters. Parameters for starch kinetics are adapted from (9, 33, 34).

| Quantity | Symbol | Value | Units |
| --- | --- | --- | --- |
| Concentration | $c$ | - | $\text{mol m}^{-3}$ |
| Effective diffusion | $\omega RT$ | - | $\text{m}^3 \text{s}^{-1}$ |
| Water potential | $\psi$ | - | Pa |
| Turgor pressure | $P$ | - | Pa |
| Water flow | $U$ | - | $\text{m}^3 \text{s}^{-1}$ |
| Sugar flux | $J$ | - | $\text{mol s}^{-1}$ |
| Transpiration | $E$ | - | $\text{mol s}^{-1}$ |
| Starch content | $s$ | - | $\text{kg m}^{-3}$ |
| Relative intercellular hydraulic conductivity | $\xi$ | $0 \leq \xi \leq 1$ | - |
| Reflection coefficient | $\sigma$ | $0 \leq \sigma \leq 1$ | - |
| Mesophyll concentration | $c_m$ | 500 | $\text{mol m}^{-3}$ |
| Free diffusion of sucrose | $D_{\text{free}}$ | $2.3 \times 10^{-10}$ | $\text{m}^2 \text{s}^{-1}$ |
| Gas constant | $R$ | 8.314 | $\text{m}^3 \text{Pa K}^{-1} \text{mol}^{-1}$ |
| Temperature | $T$ | 298.15 | K |
| Molar volume of water | $\bar{v}$ | $1.807 \times 10^{-5}$ | $\text{m}^3 \text{mol}^{-1}$ |
| Sucrose molar mass | $M_s$ | 0.34 | $\text{kg mol}^{-1}$ |
| Starch kinetics |  |  |  |
| Michaelis constant | $K_m$ | 500 | $\text{mol m}^{-3}$ |
| Maximum velocity | $V_{\text{max}}$ | 5 | $\text{mol m}^{-3} \text{s}^{-1}$ |
| Hydrolysis rate constant | $k_h$ | 0.1 | $\text{s}^{-1}$ |

**Fig. 2.**
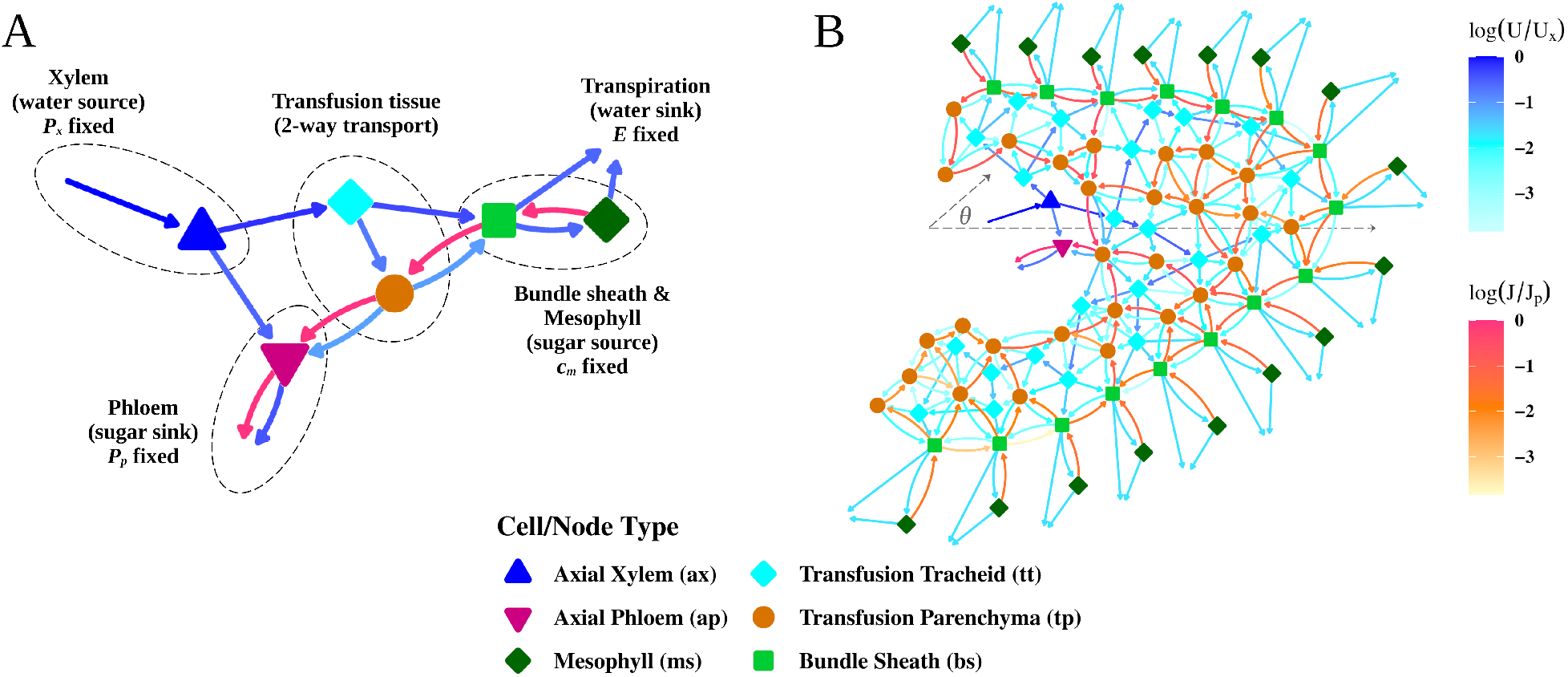
Diagrams of the steady-state solutions of the transfusion tissue network model. Node shape and color correspond to cell type. The arrows indicate water (cyan to blue) and sugar (yellow to red) flows between cells, normalized to the flows through the axial xylem and axial phloem, respectively. (A) The fundamental network model, with a summary of boundary conditions. (B) A representative network constructed from a *P. pinea* needle. The polar angle *θ* roughly describes whether a cell is towards the adaxial (*θ* > 0) or abaxial (*θ* < 0) side of the needle. An overlay of this network onto its original μCT image (from Figure 1B) is presented in the SI.

#### Governing equations

Between two nodes *i* and *j*, the water flow *U*_*i j*_ is described by the Kedem-Katchalsky equation for flow across a membrane (35):

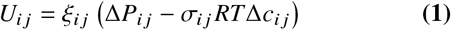

where *U*_*i j*_ indicates movement of water from node *j* into node *i* such that *U*_*i j*_ = − *U* _*ji*_. Equation 1 describes the driving force for water flow as the difference in water potentials, multiplied by a hydraulic conductivity ξ_*i j*_. The water potential is comprised of a pressure component (Δ*p*_*i j*_) and an osmotic component (−*σ*_*i j*_ *RT*Δ*c*_*i j*_), with *R* as the ideal gas constant, *T* the temperature, and *σ*_*i j*_ the reflection coefficient (discussed below). The conductivity ξ_*i j*_ is an edge property that is defined by the identities of nodes *i* and *j* and is symmetric (ξ_*i j*_ = ξ _*ji*_). Between tracheary elements (ax, tt), bordered pits mediate the flow between cells, offering low resistance and, thus, high conductivity for water flow. Between two living cells (ap, tp, bs, ms), water moves either through membrane-bound selective protein channels known as aquaporins or through plasmodesmata nanochannels, which provide cytoplasmic continuity between cells. Between a living and a tracheary cell, water travels exclusively through aquaporins embedded in the membrane of the living cell and pits of the tracheary cell. Flow through aquaporins and plasmodesmata encounters higher resistance compared to flow through the large bordered pits. Within the model, we normalize all conductivities to the conductivity through the bordered pits, such that ξ_bp_ = 1, and explore the parameter space for the conductivities across interfaces mediated by only aquaporins (ξ_aq_) and by both plasmodesmata and aquaporins (ξ_pd_). Table 2 summarizes these relationships.

**Table 2.**
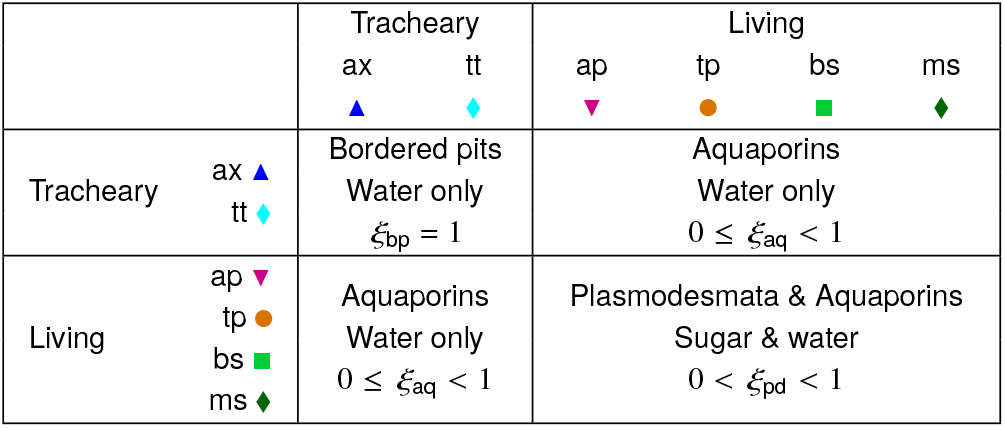
Summary of the relative intercellular conductivities, ξ, for each cell pair.

The reflection coefficient *σ*_*i j*_ (0 *≤ σ*_*i j*_ *≤* 1) is a symmetric quantity that describes the permeability of solutes (i.e., sugar) across an intercellular interface and, thus, the strength of osmotic effects. This coefficient remains undefined between tracheary elements (ax, tt), since it is only relevant for connections involving living cells (*σ*_pd_, *σ*_aq_). When *σ*_*i j*_ = 1, the membrane is impermeable to the solute, so a gradient in the osmotic pressure can be fully established. When *σ*_*i j*_ = 0, the solute can freely pass through the interface, and its flow is directly coupled to the water flow. For connections between a living and a tracheary cell, which are mediated by aquaporins, *σ*_*i j*_ = *σ*_aq_ = 1. The reflection coefficient between two living cells, *σ*_pd_, approaches 0 but can be varied within the model to bias the relative abundance and accessibility of aquaporins (higher *σ*_*i j*_) versus plasmodesmata (lower *σ*_*i j*_), since sugar can pass through plasmodesmata but not through aquaporins.

Sugar can only flow symplasmically (via the cell lumen) through the living cells (22). As a consequence, there is no sugar flow into or through the tracheary elements, and their sugar concentration is always zero. The model assumes that sugar is transported passively without the input of metabolic energy, as is generally believed in conifers. Though sugar transporters have been found in conifers, their function within the leaves remains unclear, and we ignore their potential contributions in our model (36). The sugar flow *J*_*i j*_ is described by an advection-diffusion equation:

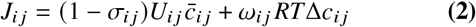

where 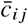 is the mean concentration of nodes *i* and *j*, a valid approximation for a low Péclet number regime. The factor *ω*_*i j*_ *RT* serves as an effective diffusion coefficient, where the solute mobility *ω*_*i j*_ is a function of the size and abundance of plasmodesmata, along with the free diffusion of sugar (37, 38).

Starch synthesis and hydrolysis are enzyme-mediated processes (39). We approximate starch (*s*_*i*_) dynamics using Michaelis-Menten enzyme kinetics:

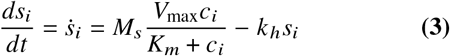

where *M*_*s*_ is the molar mass of sucrose, *V*_max_ is the maximal velocity of starch synthesis, *K*_*m*_ is the Michaelis constant indicating the concentration *c*_*i*_ at which the half-maximal velocity is achieved, and *K*_*h*_ is the rate constant for starch hydrolysis. The net change in sugar concentration in node *i* is a function of both transport and sequestration:

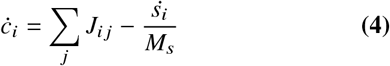

#### Model constraints and boundary conditions

For an incompressible fluid, mass conservation is ensured by continuity:

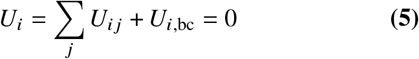

where *U*_*i*,bc_ accounts for the water flows into node *i* from its relevant boundaries. For the transfusion tracheids and transfusion parenchyma, this term is zero. The axial xylem is connected to a distant source of water at the roots with a fixed pressure. The axial phloem is connected to a distant sugar sink with a fixed positive pressure and a sugar concentration of zero. The bundle sheath and mesophyll cells are subjected to a boundary condition representing the evaporative surfaces of the leaf near the stomata, where a fixed transpiration (*E*) is imposed. *E*, given in moles per time, is a metric that can be physically measured. In the model, this corresponds to the net boundary flow from the bundle sheath and mesophyll cells, scaled by the molar volume of water,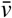.

The mesophyll is assumed to be photosynthesizing at capacity, meaning its concentration never drops below *c*_*m*_; however, in some instances, sugar is swept backward into a mesophyll cell. In this case, the concentration in the backwashed mesophyll cell is allowed to exceed *c*_*m*_ (32). Alternatively, the model can instead impose a fixed assimilation, or photosynthetic, rate at the mesophyll. At steady state, this is equivalent to imposing a fixed sugar export rate through the axial phloem, *J*_p_. The model is allowed to evolve to a steady state, with the net sugar flux through each cell *J*_*i*_ = 0 everywhere.

The model was implemented in MATLAB R2022b. The codebase, along with a description of the algorithm and parameters, is available at https://github.com/melissahmai/transfusion_tissue.

## RESULTS

### Needle architecture and stelar tissue types

The cross-sectional distribution of tissues in the needle of *Pinus pinea* revealed, from outside to inside, the epidermis with stomata, mesophyll, and bundle sheath surrounding the stele with two vascular bundles embedded in the transfusion tissue. The architecture seen in both the fixed and stained material (fluorescence and electron micrographs; Figure 1A and C) was well revealed in reconstructed cross sections from whole-needle, live μX-ray computed tomography. Here, the water-filled dead transfusion and xylem tracheids clearly contrast the living transfusion parenchyma, phloem, bundle sheath, and mesophyll cells (Figure 1B). The transfusion tissue separated the endodermis and vascular bundles completely as seen in electron micrographs. It consisted of two cell types only, transfusion tracheids and transfusion parenchyma cells, the latter of which were characterized by dark tannin vacuoles and starch grains (Figure 1C). Starch grains were also found in mesophyll and bundle sheath cells. The phloem flank was marked by a group of Strasburger cells, which aid with phloem loading, that otherwise occurred in phloem rays (Figure 1C: yellow).

### Model behavior

A representative steady-state solution of a network constructed from a *Pinus pinea* μCT image is shown in Figure 2B and behaves well with the chosen parameters. Water exits the xylem and flows towards the sites of evaporation, and sugar moves from the sites of photosynthesis to the phloem loading point for export. Redistribution of sugar is observed along the endodermis before entering the transfusion parenchyma.

Quantitative analysis of cell-level physiological metrics within the stele is also possible with the model (Figure 3). The water potential *ψ* = *P* − *RT c* describes the chemical potential of water as an energy density, with water flowing from higher to lower water potentials. Away from the xylem, *ψ* becomes increasingly negative, driving water flow from the xylem to the rest of the needle, ending in the phloem to be exported as sap, into the mesophyll for photosynthesis, or beyond the mesophyll to be evaporated near the stomata (Figure 3A). The concentration profile (Figure 3B) shows a downward gradient from the mesophyll to the phloem, consistent with the passive loading scheme in conifers.

**Fig. 3.**
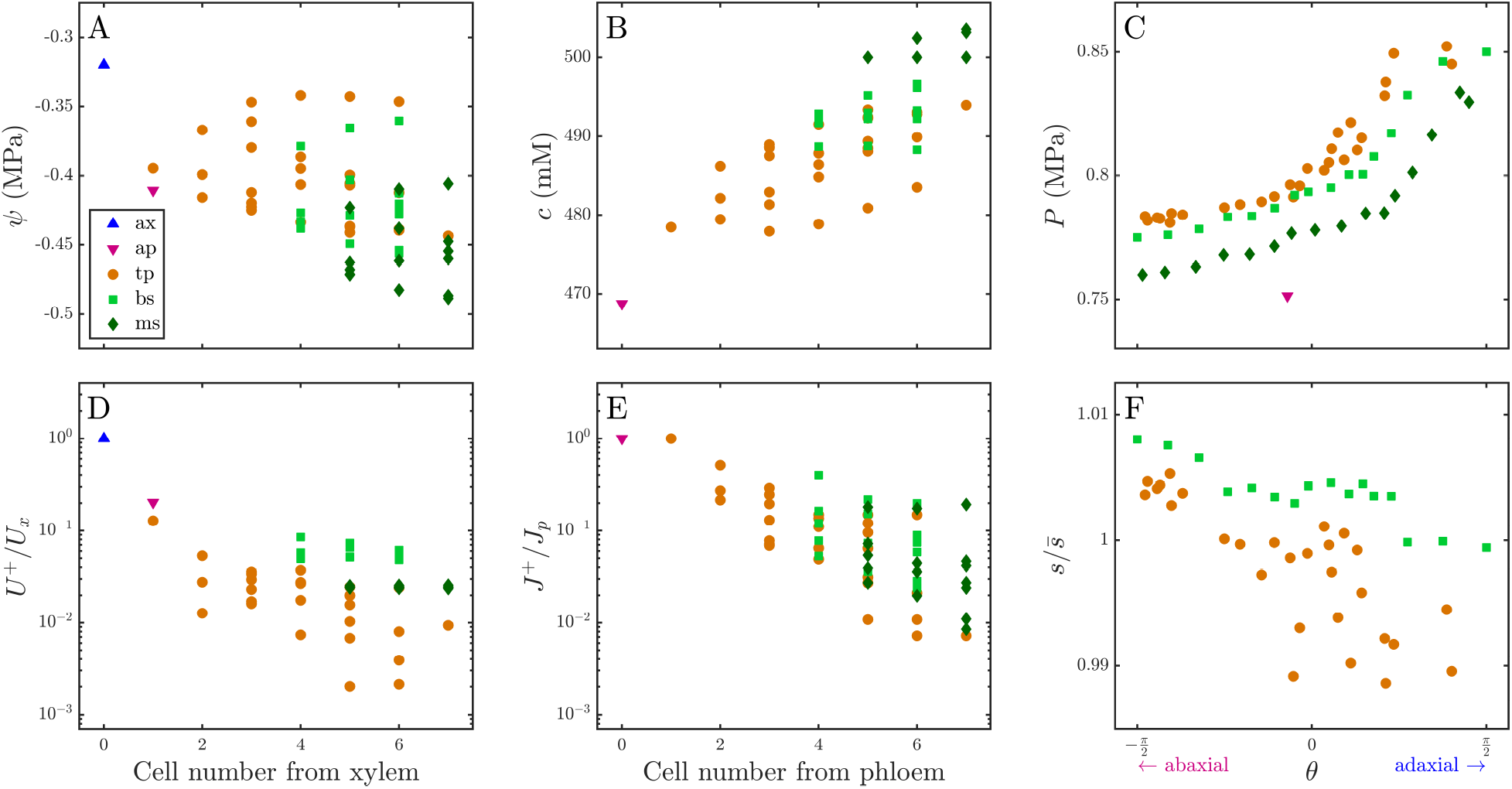
The steady-state solution of a representative *Pinus pinea* network demonstrates the effect of cell position on flow. Sugar and water flows dissipate with cell rank from the axial vasculature. Each point represents an individual cell (axial xylem (ax, dark blue triangles), axial phloem (ap, magenta downward triangles), transfusion parenchyma (tp, orange circles), bundle sheath (bs, light green squares), and mesophyll (ms, dark green diamonds)). (A) Cell water potential *ψ* in relation to cell number (i.e., shortest discrete path) from the axial xylem. (B) Sugar concentration *c* in the living cells in relation to cell number from the axial phloem. (C) The tangential distribution of the turgor pressure *P* in each cell, with *θ* < 0 corresponding to the abaxial, or phloem-adjacent, region of the needle, and *θ* > 0 corresponding to the adaxial, or xylem-adjacent region. (D-E) The positive components of the water *U*+ and sugar *J*+ flows through each cell, normalized by the total xylem water flux *U*_x_ and phloem sugar flux *J*_p_, respectively, as a function of discrete cell number from the axial flank. (F) The tangential distribution of starch content *s*, normalized by the average starch content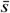.

At the interface between the endodermis and mesophyll, but elsewhere as well, the flows of sugar and water directly oppose each other. The Péclet number is a dimensionless quantity that describes the ratio of advective and diffusive transport. At low Péclet numbers, only a small concentration gradient is required for the osmotic pressure gradient to overcome the advective water flow, allowing for bidirectional transport across this interface. Figures 3D and 3E show the positive components of the total flow rates of water (*U*^+^) and sugar (*J*^+^) (such that 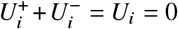, or equivalently, 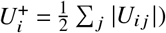, normalized by the flows through the xylem (*U*_x_) and phloem (*J*_p_), respectively for each node. For both water and sugar, the branched network structure of the transfusion tissue causes the flows to slow down by several orders of magnitude with increasing distance from the axial vasculature, allowing for the low Péclet numbers that enable this bidirectional transport observed across the endodermis.

Each cell’s tangential position within the needle can be roughly described by its polar angle *θ* with respect to the vascular bundle, based on the original μCT image the network was constructed from, with *θ* < 0 referring to abaxial (phloem-adjacent) regions, *θ* > 0 referring to adaxial (xylem-adjacent) regions, and *θ* ≈ 0 corresponding to the flank of the needle (Figure 2B). Pressures are higher on the adaxial side, where there is generally an enrichment of transfusion tracheids (Figure 3C). Increased irrigation of this region relative to the opposite side of the needle reduces the potential gradient required to fulfill transpirative demands, resulting in higher turgor pressures (less negative water potentials) for the surrounding living cells.

### Starch

Starch content was added as a storage term for sugars and can be used as an indicator for sugar concentration, as there is a simple analytical expression for the steady-state solution of Eq. (3). The inclusion of starch kinetics does not change the overall steady-state solution of pressures and sugar concentrations, regardless of the parameters chosen for Eq. (3), though the time to reach steady state and the final abundance of starch within the needle will be affected. Figure 3F shows the steady-state distribution of starch within the needle with the parameters given in Table 1. Starch content follows the sugar concentration. The model with the chosen parameters suggests a minor depression of starch content at the flanks and adaxial regions of the needle, and these differences may become more or less significant depending on the choice of parameters. While the steady-state solution is unaffected by starch kinetics, stored starches allow the needle to continue exporting sugars once photosynthesis has stopped and may contribute to continued export capacity at night.

### Bundle size

To investigate the impact of an intermediary transfusion tissue, we examine a reduced version of the model. If the stele is fully occupied by the axial vasculature without any transfusion tissue, the needle can be represented by a file of bundle sheath cells that are directly connected to the axial vasculature via a single cell at the flank (Figure 4A(i)). We use the export efficiency *J*_p_ / *E* (mol sugar per mol water) as a metric for assessing needle function. The export efficiency measures the amount of sugar loaded into the phloem relative to the amount of water lost to transpiration. Ideally, this ratio would be maximized. Increasing the radius of the stele can be represented by increasing the number of bundle sheath cells around its circumference (Figure 4A(ii)). Regardless of bundle size, flows still must focus at the vascular flank, so increasing the size of the bundle further exacerbates the existing bottleneck. This transformation diminishes the export efficiency *J*_p_ / *E*, until export becomes dysfunctional (Figure 4B). For systems with multiple bundle sheath cells, which do experience a constriction of flows at the vascular access points, including the transfusion tissue allows the water and sugar pathways to separate interior to the endodermis, allowing larger bundles to maintain a positive export efficiency. We note that transport through a model with a single bundle sheath cell is one-dimensional, so adding in a single layer of transfusion tissue into this system marginally decreases the export efficiency by introducing extra resistance along the transport pathway. In larger systems with more complicated connectivity profiles among cells, such as in real networks, degeneracy in the possible pathways further distributes these flows and can accommodate even larger steles.

**Fig. 4.**
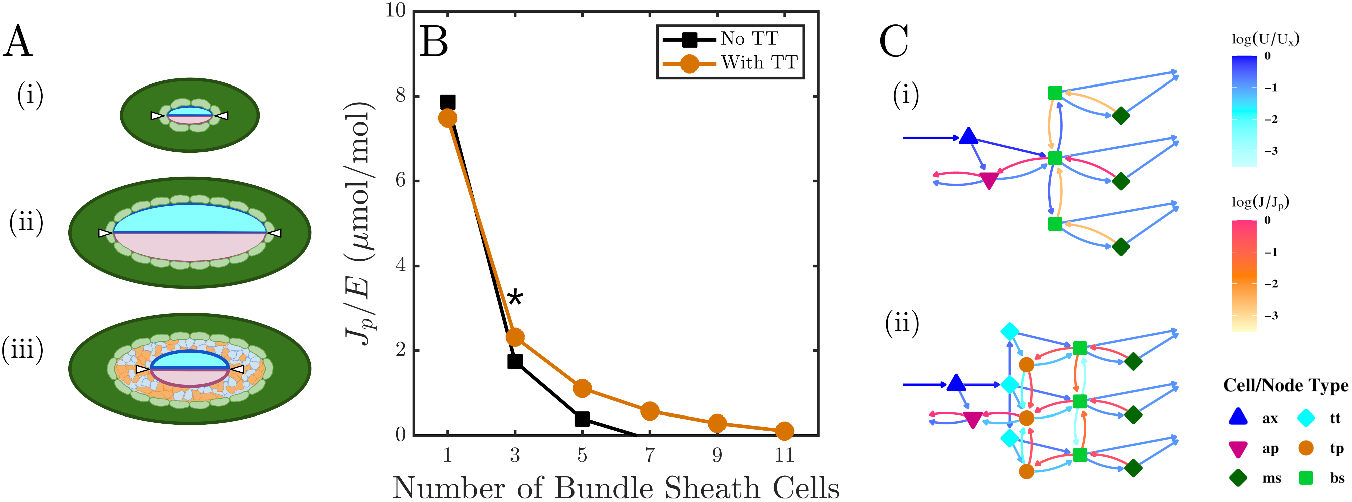
The intervening transfusion tissue alleviates bottlenecking issues. (A) Cartoons of a hypothetical stele composed of only primary vascular tissue, xylem (blue) and phloem (magenta) (i). A larger stele is circumscribed by more bundle sheath cells (pale green) (ii). However, loading and unloading is still limited to the focal points (white arrows). Transfusion tissue can be inserted into the stele (iii). (B) Increasing the size of the stele with only axial vasculature results in a rapid loss of export efficiency, *J*_p_ /*E* (black squares). Including a single layer of transfusion tissue (orange circles) bolsters export for larger steles. Diagrams of the networks marked by (*) are shown in (C), with the same symbols and colors as Fig. 2.

### Interfacial hydraulics

*Pinus*-type transfusion tissue exhibits notable compositional and spatial heterogeneity in its partitioning between transfusion tracheids and transfusion parenchyma. To investigate this, two regimes can be considered (Figure 5A). We compare (i) a fully parenchymatous tissue with (ii) a heterogeneous case, similar to what is seen naturally. In the heterogeneous model (ii), the transfusion tracheid and parenchyma systems are both in contact with the endodermis but also highly interdigitated with each other, and aquaporins in the membranes of the transfusion parenchyma along their interface allow for water to flow between them. This water flow is dictated by the difference in the water potential, which includes hydrostatic and osmotic pressure terms. By shifting the model between this configuration and a fully parenchymatous one (i), we can test the importance of physically separating the net water and sugar currents on the export efficiency *J*_p_/*E*.

**Fig. 5.**
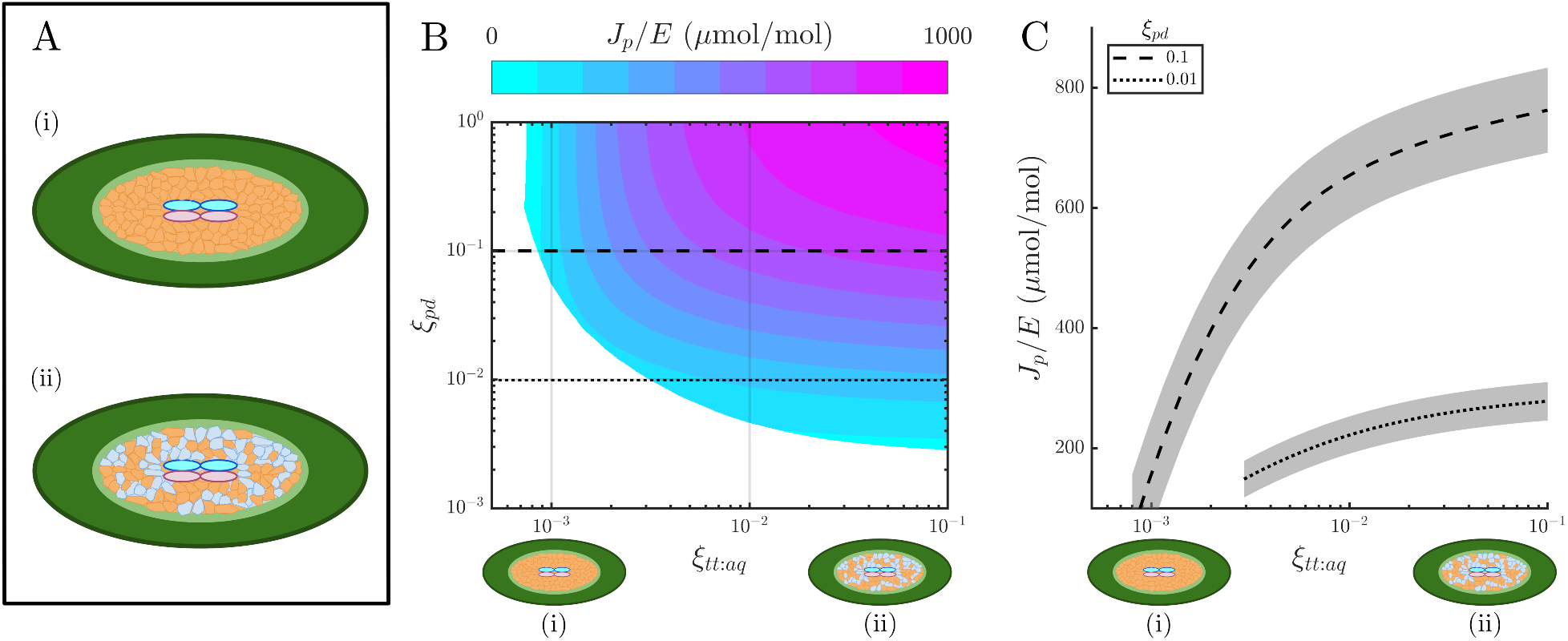
The separation of water and sugar flows close to the vascular access points is crucial for needle function. (A) Conceptual schematics for hypothetical configurations of the stelar tissue. (i) A fully parenchymatous stelar tissue. (ii) A well-connected, heterogeneous transfusion tissue, with both the transfusion tracheid and transfusion parenchyma systems in contact with each other and with the endodermis. (B) Landscape of the export efficiency *J*_p_ / *E* (μmol sugar per mol water) between configurations (i) and (ii) by varying the aquaporin-mediated conductivity ξ_tt:aq_ between transfusion tracheids and both transfusion parenchyma and bundle sheath cells, relative to the plasmodesmata-mediated hydraulic conductivity between living cells ξ_pd_. The landscape is averaged over 66 network configurations. Unshaded regions correspond to the parameter space which leads to negative sugar export and/or a loss of turgor. (C) Traces of the export efficiency landscape shown in (B) at ξ_pd_ = 0.1 (dashed) and 0.01 (dotted). Shading refers to the standard deviation across all 66 configurations.

This can be accomplished by altering *all* osmotic interfaces of the transfusion tracheids, which include the tracheid-to-parenchyma (ξ_tt:tp_) and tracheid-to-bundle sheath (ξ_tt:bs_) conductivities. Figure 5B shows the effect of this concurrent attenuation of both ξ_tt:tp_ and ξ_tt:bs_, which together we define as the transfusion tracheids’ aquaporin-mediated conductivity, ξ_tt:aq_, on the export efficiency over all 66 constructed networks. It is important to note that only the conductivity of aquaporin-mediated interfaces of the transfusion tracheids are affected, while the conductivities between the axial xylem and either the axial phloem or adjacent transfusion parenchyma (ξ_ax:aq_) remain constant. The landscape is further explored by sweeping over values for the plasmodemata-mediated hydraulic conductivity among the living cells, ξ_pd_. As ξ_tt:aq_ → 0 severs the connections between transfusion tracheids and their neighboring transfusion parenchyma or bundle sheath cells, the transfusion tracheid system becomes essentially nonfunctional or nonexistent (Figure 5A(i)). Instead, water flows from the xylem directly into the transfusion parenchyma by the vascular flank and travels through the cell file into the endodermis, counter to the sugar current. Across the whole landscape, decreasing ξ_tt:aq_ restricts export efficiency, leading to lower assimilation relative to transpiration and eventually leading to total export dysfunction (Figure 5B,C). As transfusion tracheids are removed from the system with decreasing ξ_tt:aq_, water is forced to travel via the living tissue, frustrating the opposing flow of sugar and encountering greater resistance to irrigating the photosynthetic tissue. At this limit, the needle dries out and loses turgor.

We can further probe the interfacial hydraulics of the transfusion tissue by treating the transfusion tracheid system’s interactions with the transfusion parenchyma and endodermis separately. The large interfacial contact area between the transfusion tracheid and transfusion parenchyma systems suggests that hydraulic interactions between them may be important for function (11). An internally disconnected system hydraulically separates the transfusion tracheids from the transfusion parenchyma, but leaves the endodermis in direct contact with the transfusion tracheids. This could be conceived as a spatial separation and complete hydraulic disconnection of the two systems (Figure S2A(ii)). For a more nuanced approach, this can be studied, as before, as an attenuation of water flow by decreasing aquaporin abundance at the transfusion tracheid:transfusion parenchyma interface by decreasing ξ_tt:tp_ while keeping the tracheid:bundle sheath conductivity, ξ_tt:bs_, constant. Across a large range of ξ_pd_, varying ξ_tt:tp_ shows no appreciable effect (Figure S2B), suggesting the interdigitation of the transfusion tracheid and parenchyma systems may be more of a developmental consequence than a critical feature for the internal irrigation of the transfusion tissue.

If, alternatively, the interdigitation is a mere morphological consequence that allows both the transfusion tracheids and the transfusion parenchyma to evenly service the entire endodermis, we can instead study the importance of the direct irrigation of the endodermis via the transfusion tracheids (Figure S2A(iii)). The hydraulic conductivity across this interface, ξ_tt:bs_ can be restricted while keeping ξ_tt:tp_ unaffected. Even though water flow is limited directly between the transfusion tracheids and bundle sheath cells, water can still enter the endodermis via the transfusion parenchyma. We do not study the case in which the transfusion parenchyma lose contact with the endodermis; since sugar can only flow through the transfusion parenchyma, severing its connection to the endodermis will obviously cause dysfunction. Altering these hydraulic connections between the transfusion tracheids and the bundle sheath cells also has no significant effect on function, as reported by the export efficiency *J*_p_/*E* (Figure S2C). This modification does little to change the behavior of the needle, since water can still enter the endodermis via adjacent transfusion parenchyma cells, which can draw water from the transfusion tracheids. In effect, this only extends the path length of the water flow by one cell.

## DISCUSSION

Throughout the literature, several hypotheses have been developed about the function of the transfusion tissue. These functions are not necessarily mutually exclusive; in fact, this tissue likely has multiple roles. For example, in addition to buffering sugar flow with starches, the transfusion tracheids may also serve as a “hydraulic circuit breaker” for the needle. Due to their thinner walls relative to those of the xylem, transfusion tracheids have been observed collapsing under drought, with their volume recovering upon rehydration (23, 24). This is reminiscent of drought responses in other plant taxa, such as the reversible collapse of smallest-order veins in angiosperms (40) or the complete disconnection of the vascular bundle from the photosynthetic tissue of lycopods during drought (41). As most conifers are evergreen, their needles must persist over multiple years, subject to several freeze-thaw cycles. By restricting access to water in the xylem on sunny winter days, freezing temperatures induce drought conditions for needles that are photosynthesizing and transpiring (42). Having a tissue serve as a hydraulic shock absorber protects the axial xylem from embolism, which would render it entirely nonfunctional. Yet, the fact that deciduous conifers, such as larches, or those that live in warmer climates, also have transfusion tissue suggests it provides additional advantages. Our results suggest that it separates and distributes fluid flow to overcome structural constrictions and support efficient irrigation and export.

### The model provides cell-level resolution of water and sugar flows

The transfusion tissue is a characteristic yet enigmatic feature of gymnosperm leaves and has been a subject of study since it was first described by Frank in 1864 (43). While there has been significant work on the anatomy and development of the transfusion tissue over the past century and a half, its physiology remains largely unexplored. We have developed a minimal mathematical model to resolve cell-level details of sugar and water flows through the transfusion tissue of conifer needles, a system that is difficult to probe experimentally. By exploring the parameter space, the model can overcome some deficiencies in empirical data to investigate how the structure of the transfusion tissue solves the architectural challenge of flow constriction at the focused vascular access points.

Moreover, while we have applied this model to networks built from *Pinus pinea* needles, the model is not limited to the *Pinus*-type transfusion tissue described by Ref. (21) and is widely applicable to diverse anatomical configurations of transfusion tissue. This versatility with respect to both configurations and boundary conditions underscores the model’s utility in predicting needle function under several physiological conditions. While the analysis presented here has been of steady-state solutions, the model can be adapted to understand temporal dynamics in response to environmental stimuli.

### Flow through the endodermis

The Casparian strip at the endodermis forces water to move symplasmically across the endodermis via the plasmodesmata or aquaporins (11, 12, 22). In the network model, flow can be biased such that it moves entirely through the plasmodesmata by setting the reflection coefficient *σ* to zero, and bidirectional transport is still possible. Therefore, at this interface of the endodermis, opposing flows of water and sugar may be established through the same physical channels of the plasmodesmata. The branching of flows from the vasculature toward the endodermis allows the flow velocities to be quite low at the endodermis. At these small Péclet numbers, only small concentration gradients are necessary to generate a diffusive sugar flow that opposes a convective flow (32). In real needles, water can also flow through aquaporins in the endodermis (44). When the model accounts for the presence of aquaporins (*σ* > 0), this bidirectional flow becomes even easier to accomplish, as this opens another avenue for water to move separately from the sugar current.

Though the tiered architecture of the axial vasculature has been discovered, the exact local loading scheme into the phloem along the length of each tier remains unclear. Each segment is short enough for uniform loading along its length without stagnation, but it is also conceivable for sugars to be loaded primarily at the tip of each segment (9). External to the transfusion tissue, sugars can potentially travel axially down the endodermis and enter the stele in the plane at which a new set of vascular conduits appear. Yet, plasmodesmata-rich pit fields in the endodermis have been found in abundance in the tangential direction, in contrast with sparse plasmodesmata in the axial direction. This suggests that axial transport between bundle sheath cells is limited compared to radial or tangential transport (45). Tangential redistribution of sugar along the endodermis before entering the transfusion tissue in the model is consistent with experimental observations of enrichment of plasmodesmata between bundle sheath cells. It is a reasonable assumption that the preferential tangential sugar flow in the endodermis indicates uniform axial loading of the phloem along a given segment. To interrogate this assumption, the current model may be extended to simulate concurrent axial and radial transport through networks built from three-dimensional reconstructions.

### Starch accumulation

Diurnal variations in leaf starch content have been previously studied, in both conifers (9, 46) and angiosperms (47–50). Starch synthesis during the day sequesters excess sugars to prevent excessive osmotic buildup (51). Starch accumulation cannot be used in the problem of axial stagnation, as phloem conduits lack the enzymes necessary to build or hydrolyze starches. However, the transfusion tissue may serve as a storage space to help evacuate sugars from the mesophyll and prevent end-product inhibition of photosynthesis (52). Higher sugar concentrations do lead to higher starch content in the model. When photosynthesis stops at night, starches are hydrolyzed back into soluble sugar to be exported from the needle. This increases net export capacity by obviating photosynthetic overwhelm during the day and utilizing the night to continue export.

Generally, starch accumulates in the chloroplast of the mesophyll (39), but starches have been observed in the non-photosynthetic transfusion parenchyma of conifers (Figure 1C). While the steady-state solution that we are interested in is unaffected by starch dynamics, the model can predict starch content with cellular resolution, which can be used for guiding future experimental ventures (53). We note, however, that there are likely spatial and temporal variations in enzyme expression that will affect starch dynamics within the needle.

### Partitioning of the transfusion tissue between tracheids and parenchyma

The transfusion tissue increases the size of the stele. With a larger internal radius, the annular mesophyll benefits from a larger photosynthetic surface area without increasing its own thickness, which is limited by the depth of light penetration. Additionally, a larger bundle can help to better irrigate the needle (54). However, filling up the stele entirely with axial vasculature exacerbates bottlenecking problems because loading is still limited to the vascular flanks. To overcome this problem while still increasing the size of the bundle, including the intermediary transfusion tissue allows the flow to be distributed among many smaller pathways, eventually leading to low Péclet numbers at the endodermis, where the sugar and water flows directly oppose each other.

While building transfusion tissue to increase the size of the stele is favorable, the question becomes the partitioning of that space between transfusion tracheids and transfusion parenchyma. This can be simulated by reducing the hydraulic connection of the transfusion tracheids with both the transfusion parenchyma and the bundle sheath cells of the endodermis. Attenuating these conductivities forces water to travel through the parenchyma and decimates the ability of the needle to export sugar by depressing the pressures within the living cells. On the other hand, increasing the internal irrigation of the transfusion tissue provides diminishing returns, suggesting that a balance of water-conducting tracheids and sugar-conducting parenchyma is an optimal composition for the transfusion tissue.

Extending the xylem via the transfusion tracheids towards the endodermis provides a low-resistance pathway for irrigation of the mesophyll. Moreover, extravascular irrigation is not only limited to within the stele; in *Podocarpus*, “accessory transfusion tracheids” extend the reach of the vasculature further into the mesophyll (16, 55). Yet, bringing the tracheids in contact with the endodermis does not seem to be the only advantage of the transfusion tissue’s composition. Removing these direct connections effects little change, as long as the adjacent transfusion parenchyma cells are able to mediate the movement of water at that interface. By also severing the connection to the transfusion parenchyma, we find that the separation of flows within the transfusion tissue is critical for function. In the context of a bottlenecking problem, this becomes most important in regions close to the vascular focusing point. This issue is less severe further from this point, as flows slow down enough for diffusive currents to overcome advective ones. Ref. (54) reports that, for angiosperms, which typically have much greater vein density, both the extravascular sugar and water currents have an effectively one-dimensional velocity profile. In other words, the flows move with a low, relatively constant velocity along their entire path, as opposed to focusing into or spreading out from a large advective velocity close to the vascular access points, as in gymnosperms. While a spatial separation of the two flows is not necessary for angiosperms, gymnosperms do benefit from this strategy to manage the consequences of their tiered axial vasculature.

The long, narrow geometry of conifer needles requires an architectural solution to efficiently traffic water and sugar along the leaf. To prevent axial stagnation, the axial vasculature develops tiers that segment the needle into separate, discrete loading zones. Consequently, only the outermost conduits of each segment are accessible for loading and unloading, forcing all radial water and sugar flows to focus from the entire needle circumference toward two points at the flanks. The transfusion tissue immediately separates the water and sugar pathways to reduce resistance between the vasculature and the endodermis. While the Casparian strip prevents the separation of water and sugar flows through the endodermis, the branched structure of the transfusion tissue dissipates flow velocities so that the advection of water and diffusion of sugar may directly oppose each other at that interface without frustrating sugar export.

## ACKNOWLEDGMENTS

This work was supported by the Independent Research Fund Denmark in the grant Multimodal imaging and modelling of vascular flows in leaves (Grant no. 9040-00349B) and by the National Science Foundation through the Harvard University Materials Research Science and Engineering Center (DMR-2011754). We acknowledge the Paul Scherrer Institut, Villigen, Switzerland, for provision of synchrotron radiation beamtime at the TOMCAT beamline X02DA of the SLS and would like to thank Dr. Margaux Schmeltz. MHM recognizes support from the Fannie and John Hertz Foundation Fellowship and the National Science Foundation Graduate Research Fellowship and thanks Sophie Everbach, Tinker Green, and Tony Rockwell for helpful discussions.

## Notes

https://github.com/melissahmai/transfusion_tissue

### Competing Interest Statement

The authors have declared no competing interest.

### Summary of Updates

Grammatical errors fixed; Figure 4 updated; Additional clarifications in the results section

