## Supplemental Information for "Relieving the transfusion tissue traffic jam: a network model of radial transport in conifer needles"

### Supporting Information Text

**Code and data availability.** The codebase for the transfusion tissue network model, along with the images and network descriptors used for the simulations are available at https://github.com/melissahmai/transfusion_tissue.

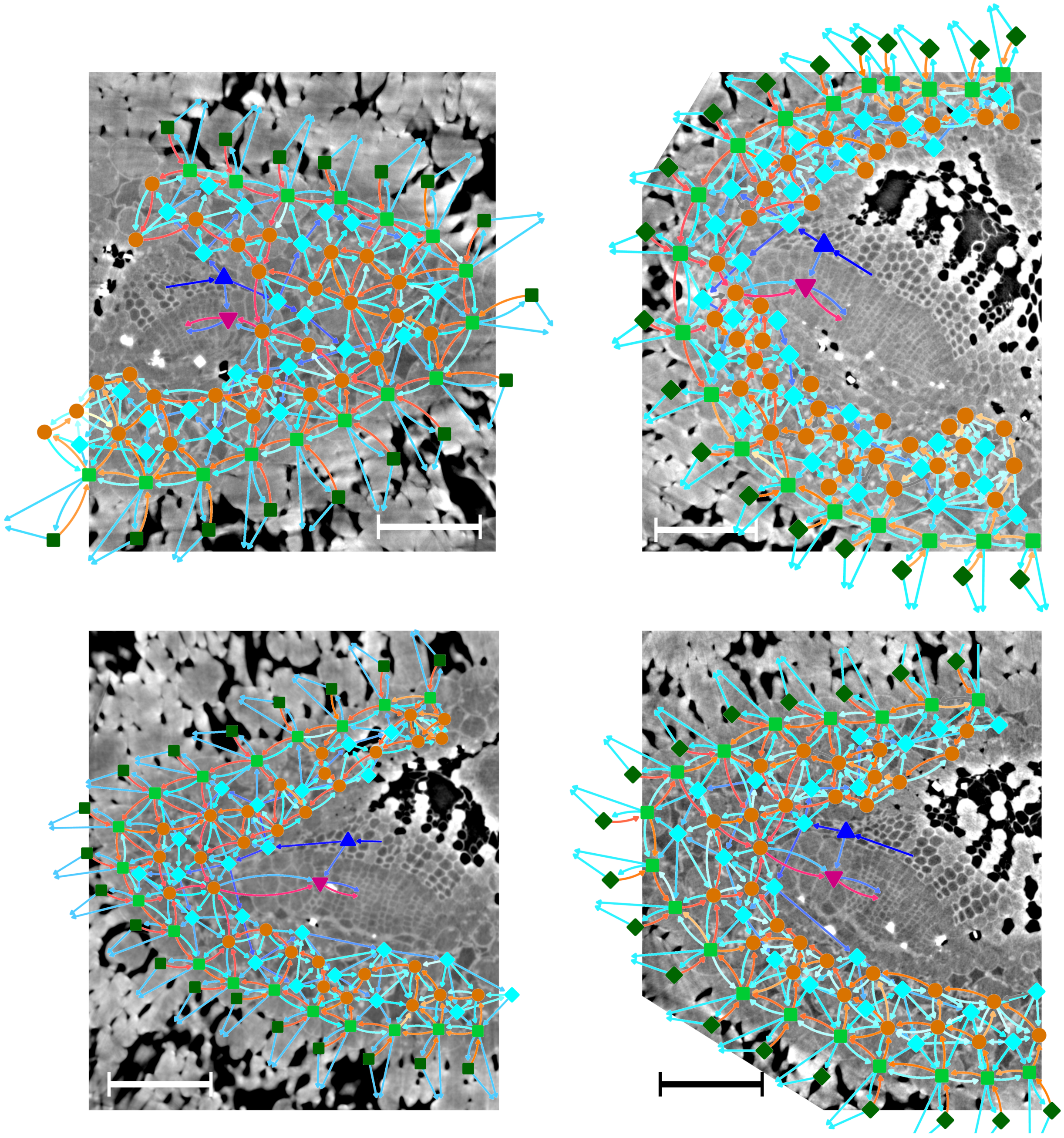

**Fig. S1.** Steady-state solutions of representative *Pinus pinea* networks overlaid on the $\mu$CT images they were constructed from. The top left network is the network discussed in the main text. Scale bars: 100 $\mu$m.

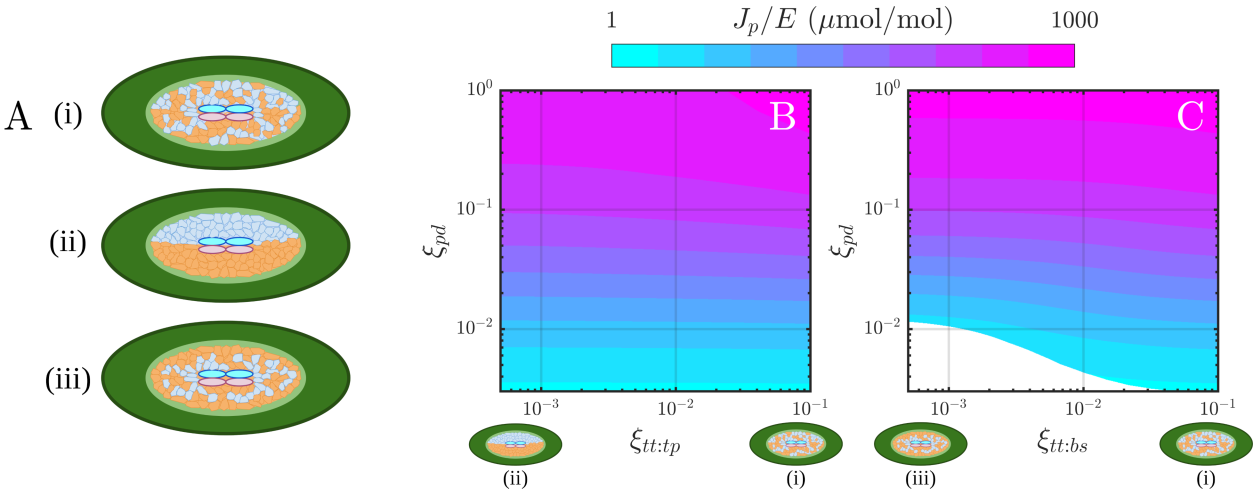


**Fig. S2.** Interfacial hydraulics. (A) Conceptual diagrams for hypothetical stelar configurations. (i) A well-connected, interdigitated case, similar to what is seen naturally. (ii) The transfusion tracheids and transfusion parenchyma are hydraulically isolated from each other, but both are still in contact with the endodermis ($\xi_{\mathrm{tt:tp}} \to 0$) (iii) The transfusion tracheids and parenchyma are hydraulically connected with each other, but the transfusion tracheids are unable to access the endodermis ($\xi_{\mathrm{tt:bs}} \to 0$). (B and C) Landscapes of the export efficiency $J_\mathrm{p}/E$ ($\mu$mol sugar per mol water) as the stelar configuration shifts from (i) to (ii) or (iii) by attenuating the relative conductivity of the transfusion tracheids with the transfusion parenchyma ($\xi_{\mathrm{tt:tp}}$, B) and bundle sheath cells ($\xi_{\mathrm{tt:bs}}$, C), respectively.
